# Accounting for location uncertainty in azimuthal telemetry data improves ecological inference

**DOI:** 10.1101/281584

**Authors:** Brian D. Gerber, Mevin B. Hooten, Christopher P. Peck, Mindy B. Rice, James H. Gammonley, Anthony D. Apa, Amy J. Davis

## Abstract

Characterizing animal space use is critical to understand ecological relationships. Despite many decades of using radio-telemetry to track animals and make spatial inference, there are few statistical options to handle these unique data and no synthetic framework for modeling animal location uncertainty and accounting for it in ecological models. We describe a novel azimuthal telemetry model (ATM) to account for azimuthal uncertainty with covariates and propagate location uncertainty into ecological models. We evaluate the ATM with commonly used estimators in several study design scenarios using simulation. We also provide illustrative empirical examples, demonstrating the impact of ignoring location uncertainty within home range and resource selection analyses. We found the ATM to have good performance and the only model that has appropriate measures of coverage. Ignoring animal location uncertainty when estimating resource selection or home ranges can have pernicious effects on ecological inference. We demonstrate that home range estimates can be overly confident and conservative when ignoring location uncertainty and resource selection coefficients can lead to incorrect inference and over confidence in the magnitude of selection. Our findings and model development have important implications for interpreting historical analyses using this type of data and the future design of radio-telemetry studies.

## Introduction

Understanding animal space-use and its implications for population and community dynamics is a central component of ecology and conservation biology. The need to understand animal spatial relationships has led to the increasing refinement and utility of telemetry devices (Millspaugh et al. 2001). Traditional telemetry data were solely collected using VHF (“very high frequency”) radio signals to track individual animals with radio tags; VHF radio-telemetry started around the mid-1960s and is still often employed. These data are collected by observers recording azimuths in the direction of the radio signal from known locations. Modern telemetry data are often collected using Argos satellites, aerial location finding (i.e., via fixed-winged aircraft), or the global positioning system (GPS). While newer forms of telemetry data are often collected, radio-telemetry devices are still relatively inexpensive. They also typically have low energy requirements, which allows for miniaturized and long-lasting devices to be fixed to small and volant animals for obtaining high spatial resolution data with minimal risk to incurring costs on survival and movement (Ponchon et al. 2013). More so, digital VHF is quickly becoming an important way to monitor the movements of small-bodied species at regional scales (Loring et al. 2017).

It is well recognized that spatial locations from telemetry devices are not without error and estimation uncertainty (Frair et al. 2004; Patterson et al. 2008). Observed locations contain measurement errors, or deviations between the recorded telemetry location and the true location of the animal. The magnitude of these deviations and the shape or structure of spatial location uncertainty is often specific to the type of telemetry technology (Costa et al. 2010) and the environmental conditions (Frair et al. 2004; White and Garrott 1990). Failing to account for location uncertainty can have important impacts on spatial analyses of animal resource selection (Montgomery et al. 2010), distribution (Hefley et al. 2014), and movement modeling (Hooten et al. 2017); location uncertainty may sometimes be modeled as a multivariate Gaussian process, but is often more complex (Costa et al. 2010).

Recent model developments focusing on satellite-based telemetry data (e.g., GPS, Argos) have highlighted the importance of appropriately characterizing location uncertainty and synthetically incorporating this uncertainty, using hierarchical modeling techniques, into ecological process models (e.g., RSF: Brost et al. 2015; Movement analyses: Buderman et al. 2016). Developments addressing the unique issues of azimuthal telemetry data do not exist; there have been few model developments to improve animal location estimation or uncertainty in the recent decades (Lenth 1981; Guttorp and Lockhart 1988). Standard practice is to analyze azimuthal data using a maximum likelihood estimator (MLE) or weighted MLE (M-estimators) to reduce the influence of outliers. These estimators are implemented in the software LOCATE (Nams 2000) and LOAS (Ecological Software Solutions LLC, Sacramento, California). Spatial location estimates are then commonly used in a secondary ecological model, in which the location uncertainty is ignored and possibly unreported, the magnitude of the uncertainty is used to define the scale of inference rather than the ecological question, and location estimates are often omitted (Saltz 1994; Withey et al. 2001; Montgomery et al. 2010). These approaches raise several concerns.

Foremost is that these practices degrade ecological inference by disregarding uncertainty, excluding data, or altering their scale of inference. Second, uncertainty from Lenth’s MLE or M-estimators are commonly defined using confidence ellipses based on the assumption of asymptotic normality (White and Garrott 1990). Assuming the uncertainty is strictly elliptical (e.g., multivariate Gaussian) may be overly restrictive and thus misrepresenting the true uncertainty. This is suggested from empirical evidence that 95% confidence ellipses of Lenth’s MLE or M-estimators cover the true location much less than 95% of the time (between 39% and 70%; White and Garrott 1990). There are also concerns raised by Lenth (1981) over the validity of the variance-covariance matrix of the M-estimators. Lastly, there are additional improvements that could add flexibility in how researchers approach the design of radio-telemetry studies. For example, Lenth’s estimators cannot estimate locations or a measure of uncertainty when only two azimuths are collected. It is also not uncommon for the estimator to fail with three or more azimuths, resulting in the use of a secondary estimator (i.e., a component-wise average of all azimuthal intersections) that has no measure of uncertainty or robust statistical properties.

Furthermore, it is well known that radio-signal direction can be influenced by many factors, including vegetation, terrain, animal movement, observer experience, and the distance between the observer and the animal (White and Garrott 1990; Millspaugh et al. 2001). To accommodate these factors, standard practice has been to test observers taking azimuths on known locations of a radio-signal to experimentally quantifying telemetry error. This error can then be applied to estimate location uncertainty via error polygons and confidence ellipses (Withey et al. 2001). If field trials obtain data across known influencing factors, a model can be developed to incorporate variation in telemetry error for these conditions (Pace and Weeks 1990). However, field trials will always be limited in their ability to anticipate all combinations of influential factors when collecting radio-telemetry data. Also, there are inconsistent recommendations in the literature regarding how best to estimate location uncertainty (White and Garrott 1990; i.e., Error polygons vs Lenth’s confidence ellipses). We developed an approach that accommodates pre-existing data sources, where field trials may not be available; if these data are available, it could be incorporated.

We developed hierarchical azimuthal telemetry models (ATM) that estimate animal locations with uncertainty, which can be synthetically propagated into spatial ecological models. We first describe a novel Bayesian ATM, which models azimuthal uncertainty using covariates. We evaluate the ATM and Lenth’s estimators under a variety of study designs. Model development is motivated by a telemetry study on the threatened Gunnison sage-grouse (*Centrocercus minimus*; Rice et al. 2017), which we use to setup the simulation and explore observer effects using the ATM. Second, we develop hierarchical spatial models for azimuthal data, including an RSF and home range analysis, which we fit to the Gunnison sage-grouse data; see Appendix S1 for species background information and study details. We examine how ignoring location uncertainty can affect ecological inference through these empirical examples, but also more generally by conducting an RSF simulation.

## Azimuthal Telemetry Model (ATM)

Suppose that multiple individuals (*l* = 1, *…, L*) are fitted with a radio-transmitter and are subsequently relocated on certain days (*i* = 1, *…, N*_*l*_). For each relocation, an observer records a set of azimuths (*θ*_*lij*_; *j* = 1, *…, J*_*li*_) at known locations **z**_*lij*_ *≡* (*z*_1*lij*_, *z*_2*lij*_)^′^ to estimate the individual’s spatial location, ***μ***^*li*^ *≡* (*μ*_1*li*_, *μ*_2*li*_) ^′^. We consider the observer locations as a fixed part of the study design and the azimuthal data observed with some uncertainty, which can be described by a circular probability distribution. We use the von Mises distribution and a trigonometric link function to relate the true animal location with the data,

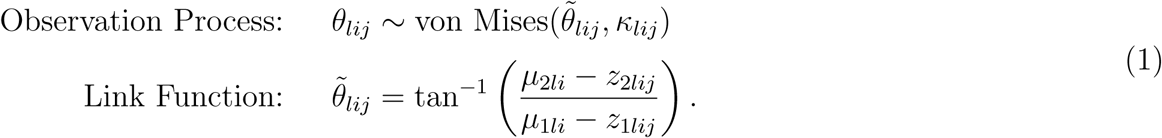

Uncertainty in the azimuthal data is controlled by the concentration parameter *κ*, in which larger values indicate less uncertainty (Appendix S2: Fig. 1), which can be modeled via covariates (e.g., observer effects; defined by the matrix **w**_*lij*_) in a hierarchical structure that accommodates unmodeled heterogeneity based on variance parameter *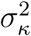*, as 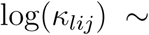 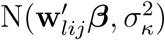. Using this framework, we can include covariates that have been hypothesized to effect azimuthal uncertainty, but have not been able to be explicitly modeled in previous studies, such as distance effects between the animal and observer, or even terrain complexity.

**Figure 1.**
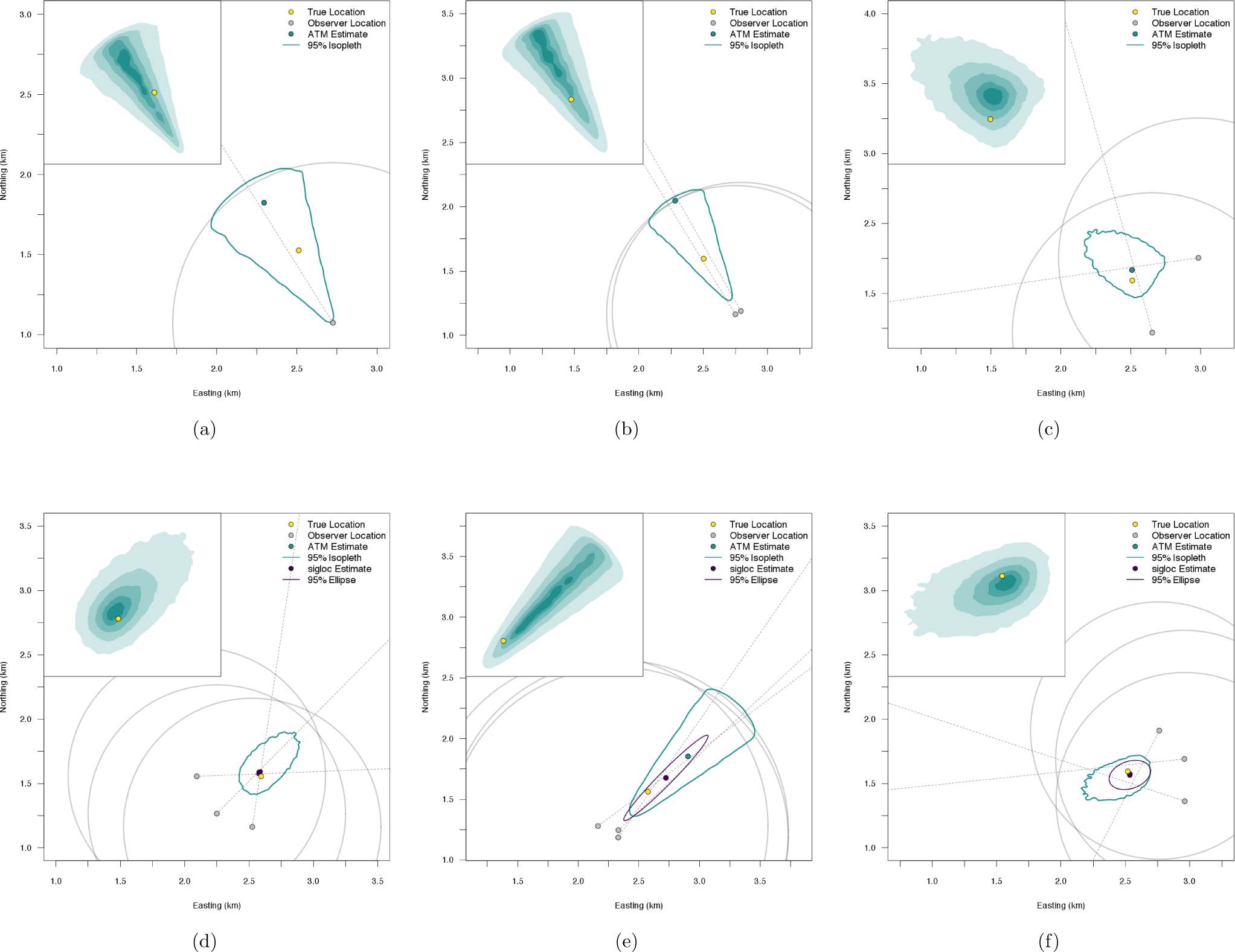
Illustrative examples of animal location estimates from the azimuthal telemetry model (ATM) and Lenth (1981) maximum likelihood estimator (*k* = 25). The union of the circles are a uniform prior probability density for the spatial location. The inset is the posterior distribution fromtheATMatisoplethsof10, 25, 50, 75, and95%. Plotswithoutasiglocestimateoruncertainty ellipse are due to estimation failure.

To complete the Bayesian model formulation, we specify priors for our unknown parameters. Commonly used priors are *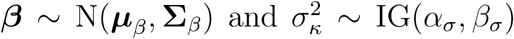*. The prior for ***μ***_*li*_ may be specified a number of ways, including multivariate Gaussian. However, to increase computational efficiency when fitting the model, it is advantageous to define an upper bound to the distance for which a telemetered individual can be detected. Otherwise, in cases where a limited number of azimuths are available or azimuths do not intersect (e.g., parallel azimuths), a multivariate Gaussian distribution will allow the uncertainty to theoretically propagate over an infinite spatial domain. In what follows, we specify a fixed maximum distance from each observer location to the animal location, using radius *r*. We also define a diffuse prior density for each spatial location as the union of all circles of the *j*^*th*^ observer location with radius *r* where **v** are coordinates (x, y) in the spatial domain,

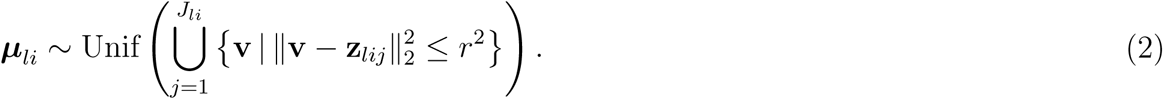

The precision of animal location estimates largely depends on the number of azimuths and whether these azimuths intersect each other. Example location estimates and associated uncertainty demonstrate the flexibility of the ATM in fitting azimuthal data with one or more intersecting or non-intersecting azimuths (Figs. 1 and Appendix S2: Fig. 2). Using Gunnison sage-grouse telemetry data from two observers, we fit the ATM to investigate possible observer differences in *κ*; the model was fit using a Markov chain Monte Carlo (MCMC) algorithm written in R (Appendix S3). We found observer one was generally more precise than observer 2 (Fig. 2). This demonstrates how we can accommodate general and specific forms of heterogeneity in *κ*, which was not previously possible with other methods.

**Figure 2.**
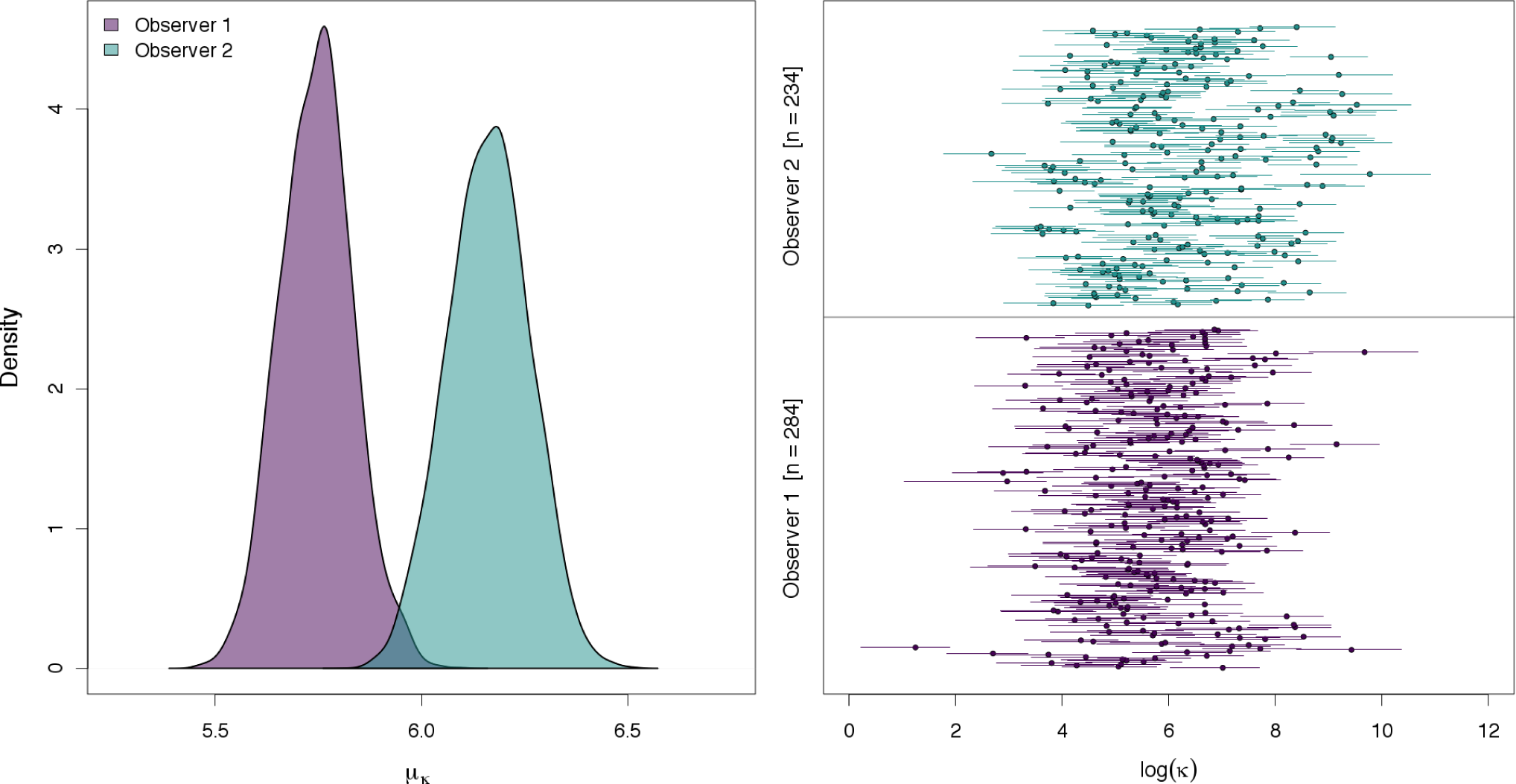
Posteriordistributionsofestimatedobservereffectsonazimuthaltelemetryuncertainty (left, *k*) andindividuallocation *k* (right; circles aremediansof theposteriordistributionandbars are 95% credible intervals) for Gunnison sage-grouse data in 2009.

## Simulation

We evaluated the performance of the ATM and Lenth’s MLE and M-estimators (Andrews and Huber) along with a simple component-wise average of intersections. We did so by simulating data under two common radio-telemetry study designs (road and encircle) and a more variable approach (random). The random design placed observers at any combination of angles from each other and to the animal location. The road design constrains observer locations to a linear feature, thus limiting the angular differences among azimuths. Lastly, the encircle design placed observer locations such that they encircled the animal location. For each design, we considered scenarios of 3 or 4 azimuths per location and moderate and high azimuth uncertainty (*κ* = 100 or 25, respectively). The distances between observer and animal locations were drawn by randomly selecting empirical distances estimated from the Gunnison sage-grouse data (Appendix 2: Fig. S3). Simulation algorithms are provided in Appendix S3 and available R code. The ATM, assuming a homogeneous *κ*, was fit using MCMC. Lenth’s MLE and M-estimators were fit using Lenth’s original algorithms (Lenth 1981; see R code). Lenth’s MLE was also fit using the R package ‘sigloc’ (Sergey 2014), which does not use the algorithm suggested by Lenth (1981), but a quasi-Newton optimization algorithm which Lenth (1981) suggested avoiding.

Across scenarios, we found that locations were typically estimated from all models and estimators, except for sigloc, which had a success rate from 52 to 99%, depending on the scenario (Table 1). The ATM and simple average of intersections always produced a location estimate. Point estimates were more accurate under the encircle study design and under moderate azimuthal uncertainty; accuracy improved 1.5 to 2.5 times with four azimuths compared to three. For all scenarios, point estimates were mostly similar among the different models and estimators. However, sigloc was less accurate than the others under the random and road designs when azimuthal uncertainty was high. The most important difference we found was that of coverage of the true value. All approaches produced relatively poor coverage (0.3 to 0.6, range) except for the ATM, which proved to be slightly below nominal coverage (*≈* 90% coverage of true value).

**Table 1.**
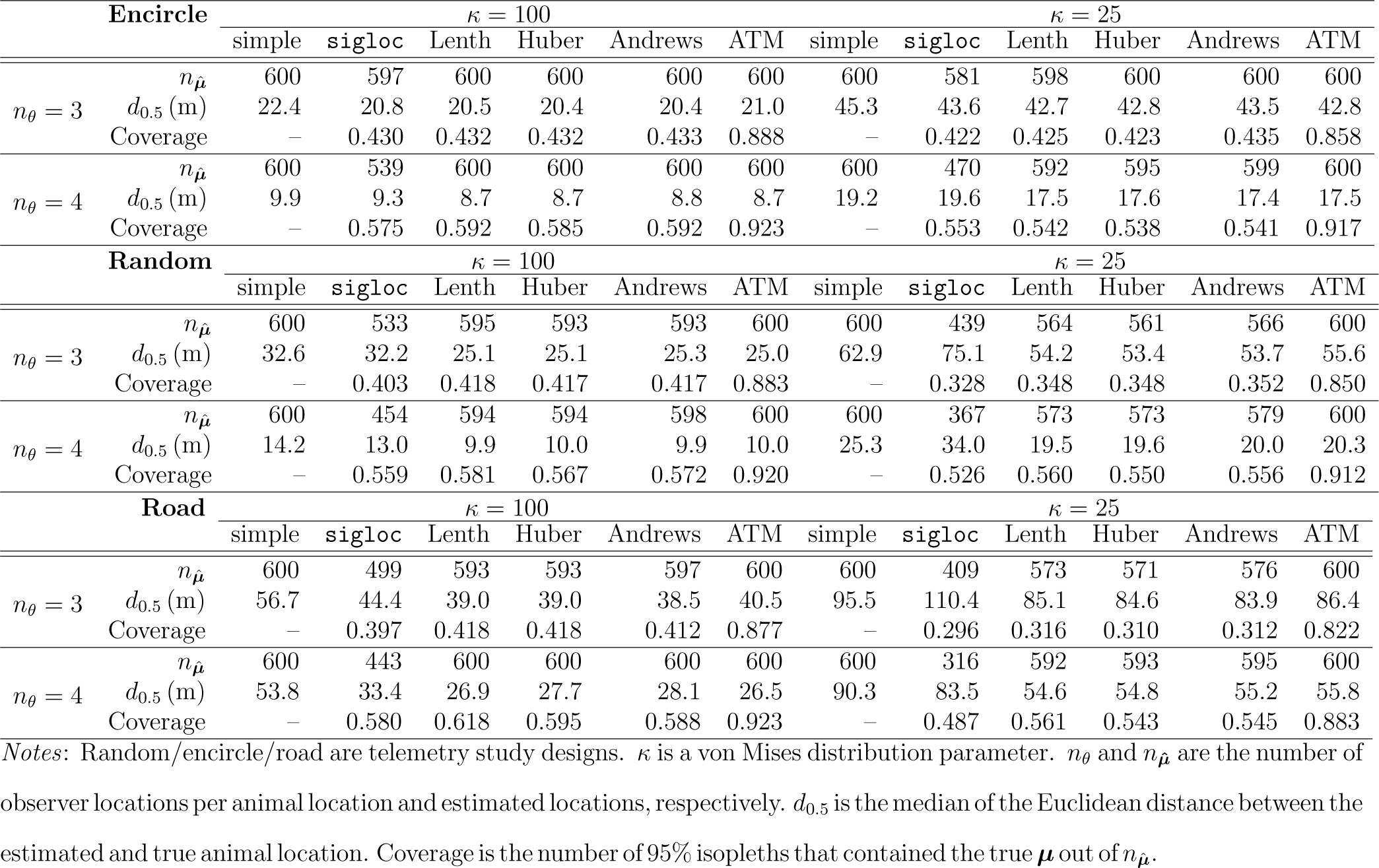
Comparing the ATM, the average azimuth intersections (simple), and Lenth’s (1981) maximum likelihood estimator (MLE; Lenth) and M-estimators (Andrews, Huber). ‘sigloc’ uses an alternative optimization for the MLE.

## Hierarchical spatial models for azimuthal data

### Resource Selection Analysis

Given our new telemetry data model, we can now analyze our estimated animal spatial locations using any ecological process model. To make inference on the relative selection of spatial resources for the population of radio-tagged individuals, we use a spatial point process, assuming independence among spatial locations (Hooten et al. 2017). Let **x** be a vector of covariates associated with location ***μ***_*li*_ and individual availability defined by the function *f*_*A*_ and availability coefficients ***θ***. Individual-level selection coefficients (***γ***) are realizations from a population-level selection process with mean and covariance (***μ***^*γ*^, **Σ**_*γ*_, respectively; Hooten et al. 2017). For multiple individuals, the hierarchical RSF model is specified as,

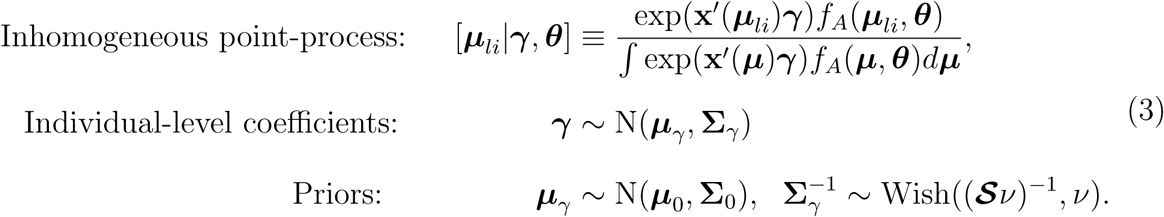

We fit the ATM-RSF model to each of a subset (six individuals) of Gunnison sage-grouse during the summer months (16 July to 30 September, from 2005 to 2009). We use these individuals as exemplars to compare estimated regression coefficients from the ATM-RSF with estimates from the same RSF, but we assumed location estimates from Lenth’s MLE are known without uncertainty. We include six common spatial variables used in RSF analyses for Gunnison sage-grouse (Appendix S1; Rice et al. 2017): road density, distance to highway, distance to wetlands, distance to conservation easements, elevation, and vegetation classification (i.e., grassland, agriculture). In addition to including both categorical and continuous spatial covariates, the variables include a highly variable topographic variable and more smoothly continuous measures of distance to features. The structure of each type and how variable values are from neighboring locations could differently impact RSF inference by the scale and shape of animal location uncertainties (Montgomery et al. 2011).

We assumed uniform spatial availability for an individual animal. To demonstrate the differences in inference, we defined the spatial extent of the availability in two ways: 1) using the convex hull of all locations (***μ***_*li*_) and 2) defining a larger study area region. The first focuses on a second-order selection process within an individual’s area of use (Johnson 1980), while the second is a first-order selection process within the broader landscape. In addition to producing fundamentally different inference for resource selection, the location uncertainty affects each differently. For the study area region, resource selection is subject to only location uncertainty, whereas for convex hull availability, resource selection is subject to both location and availability uncertainty.

As expected, resource selection depends on how we measure resource availability and whether we include location uncertainty (Fig. 3a, Appendix S4: Figs. 1-5). For example, road density is negatively selected at the study area region, but is slightly positively selected at the home range (Fig. 3a). Additionally, elevation is positively selected at the study area region, but is selected in proportion to availability (i.e., 95% credible interval includes zero) at the home range level. We found that properly accounting for location uncertainty does not always increase parameter uncertainty (Fig. 3a, Appendix S4: Figs. 1-5). Across individuals, we found the categorical vegetation variables were most affected by incorporating location uncertainty, such that including location uncertainty shifted the probability density more negative, even changing the inference and interpretation of the amount of evidence for selection of grasslands to avoidance of grasslands under the study area availability definition. The continuous variables were largely not affected when including location uncertainty, likely due to small location uncertainty relative to the adjacent spatial variability in covariate values. Lastly, an advantage of the hierarchical ATM-RSF model is that selection coefficients can inform the location estimation to where individuals were and were not likely to be on the landscape, thus reducing location uncertainty (Fig. 3b).

**Figure 3.**
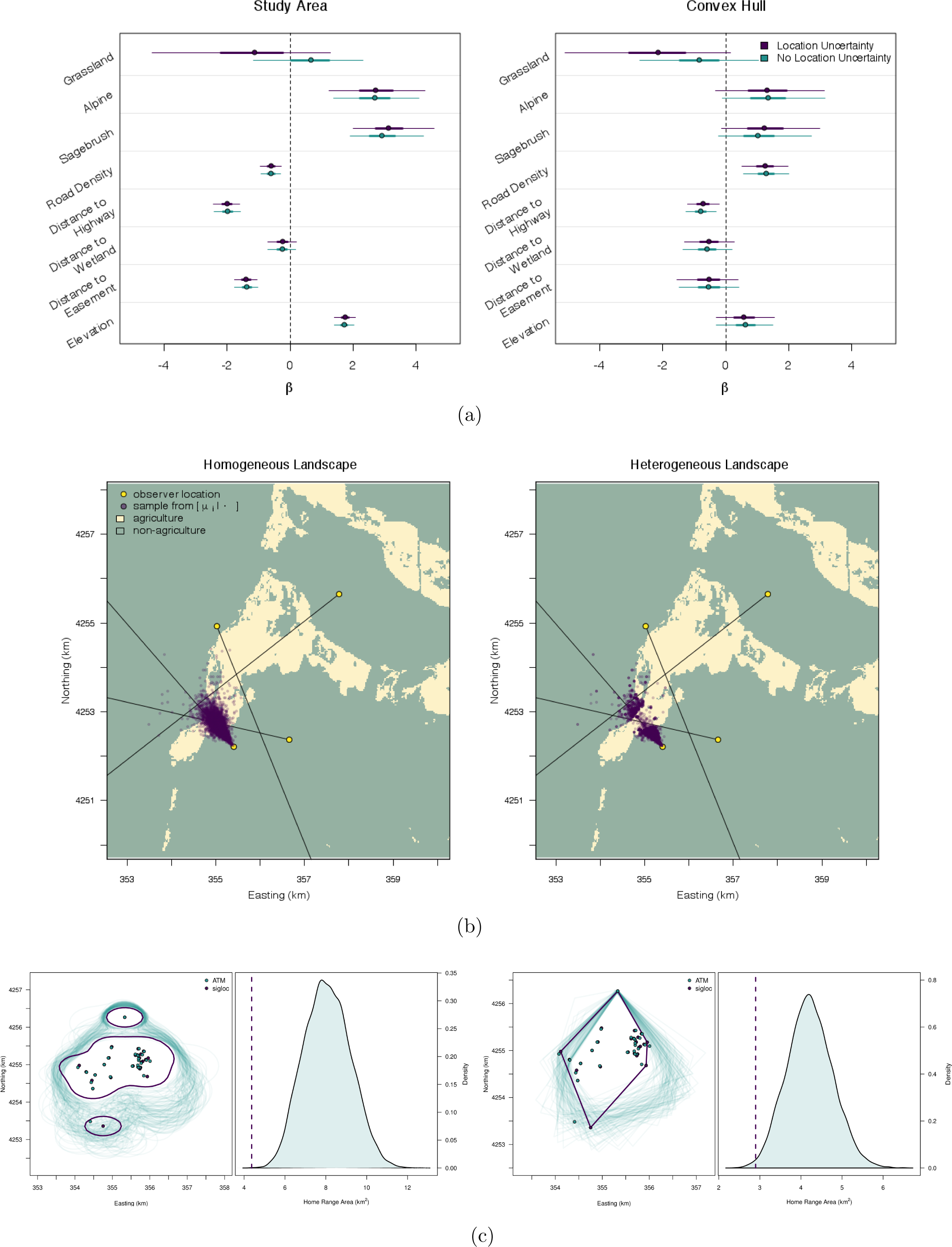
a) Resource selection coefficients for Gunnison sage-grouse; points are posterior medians, thick and thin lines are 50% and 95% credible intervals, respectively. b) Posterior samples of Gunnison sage-grouse data fit with the ATM-RSF (heterogenous landscape) and only the ATM (homogenous landscape). c) Home range distribution and area estimates for an individual Gunnison sage-grouse via kernel estimation (left) and convex hull (right) where spatial location uncertainty is incorporated via the ATM or ignored using Lenth (1981) estimation. The vertical line isthehomerangeareaestimatewhenusing Lenth(1981) estimationandlocationuncertainty is ignored.

**Figure 5.**
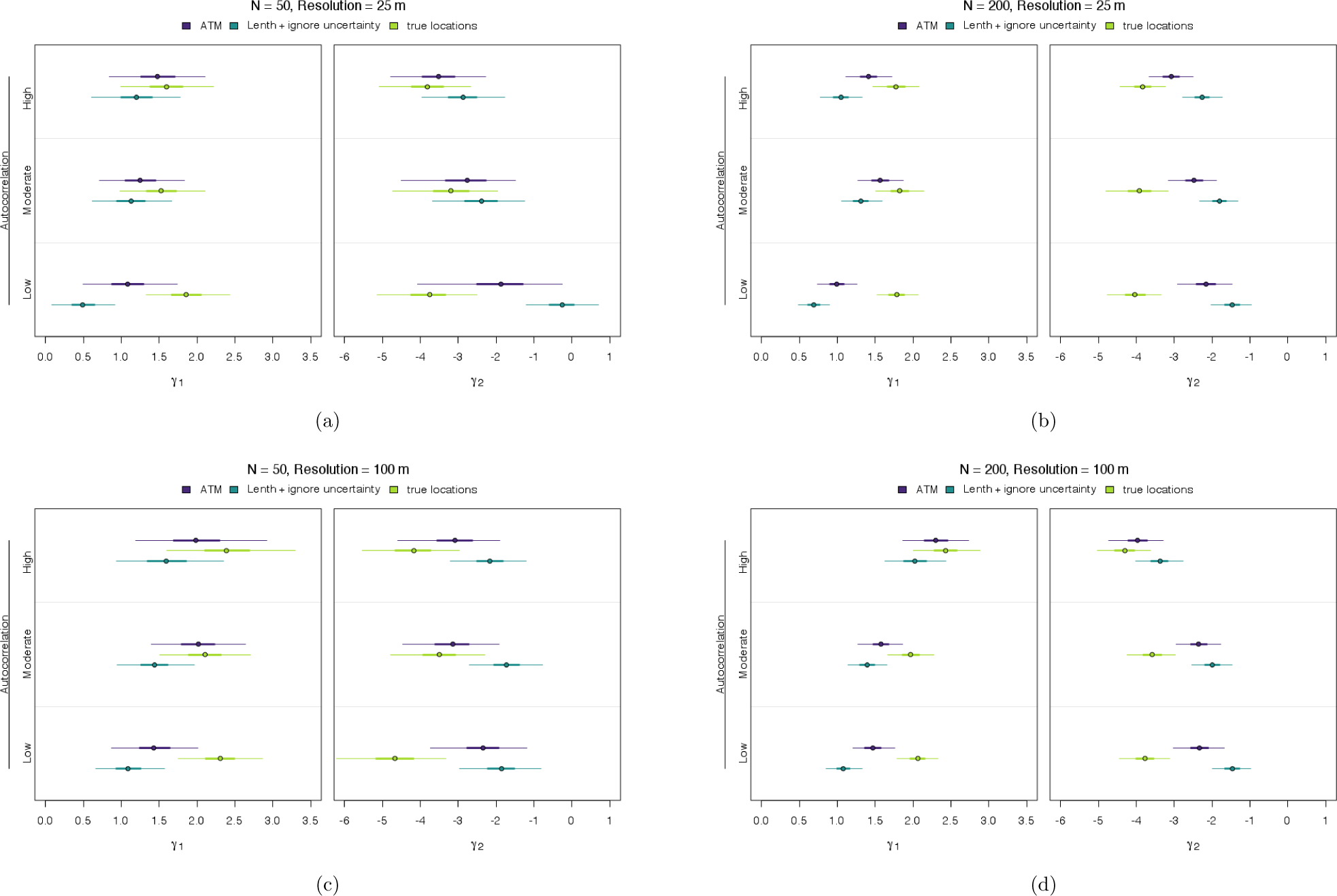
Simulation results of coefficient estimates from an RSF that incorporates location un-certaintyviathe ATM, Lenth’s(1981) maximumlikelihoodestimateswherelocationuncertainty is ignored, and when the true spatial locations are known with complete certainty. Coefficient point estimates correspond to a continuous and categorical variable (*γ*_1_, *γ*_2_, respectively) under low to high autocorrelation. Thick and thin lines are 50 and 95% credible intervals,respectively. The top row (a, b) used high spatial resolution covariates (25 m) and the bottom row (c, d) used low spatial resolution covariates (100 m). The columns differ in the size of the simulated dataset: 50 or 200 locations.

For a more general understanding, we conducted a simulation to explore the connection among location uncertainty, covariate spatial heterogeneity, and ecological inference in RSF analyses. Previous work has demonstrated this to be the case (Montgomery et al. 2011); we further this understanding by examining how varying levels of spatial autocorrelation of a continuous and categorical covariate at different sample sizes and spatial resolution effects RSF coefficients when incorporating and ignoring location uncertainty, compared to knowing the true locations. Specifically, we simulated animal location data (*N*_*locations*_ = 50, 200) that coincide with covariate values of low, moderate, and high spatial autocorrelation, defined using a Gaussian random field (covariates at 25 m or 100 m resolution; Appendix S5). Observations were three azimuths per location, simulated under a random design (Appendix S3), with moderate azimuthal uncertainty (*κ* = 50). We fit these data with 1) the ATM-RSF, and 2) a typical RSF model that used location estimates from Lenth’s (1981) MLE, ignoring location uncertainty. We compare coefficient estimates from these approaches across simulations with that of fitting an RSF where the true locations are known, providing a reference to the best case scenario for these data.

We found that differences in regression coefficients among approaches increased as spatial autocorrelation in the covariate value decreased (thus, higher spatial heterogeneity; Fig. 4). This was the case for both sample sizes and spatial resolutions, however, there was much greater uncertainty with datasets of 50 locations, compared to that of 200. Under all conditions, accounting for location uncertainty results in intervals overlapping the credible interval based on true locations to a higher degree compared to ignoring location uncertainty (Fig. 4). The difference between the ATM-RSF coefficients and those when an RSF model is fit with the known locations reflect our findings that the ATM does not always estimate loca-tions with the highest posterior density centered on the true location (with high uncertainty in *κ*; Table 1); instead, the true location is often captured in the 95% posterior isopleth. While we found that incorporating location uncertainty improves our inference about RSF regression coefficients, compared to ignoring location uncertainty, further improvement can be gained by decreasing our azimuthal uncertainty (*κ*) or increasing our certainty in animal location by taking many more azimuths (Table 1). Lastly, we found little difference among coefficients due to the spatial resolution of covariates (25 m vs 100 m); the most pronounced change was that covariates with high spatial autocorrelation and a lower resolution (100 m) led to similar coefficient estimates regardless of location uncertainty compared to those with high resolution covariates (25 m; only at the high sample size of N = 200).

### Home range

Another common use of telemetry data is to estimate the home range area of individuals. This has often been done using a convex hull or non-parametric kernel density estimation (Hooten et al. 2017). We can propagate location uncertainty using the ATM by treating the home range estimate as a derived quantity. For a given individual that was relocated *n* times within a season, we can estimate their seasonal home range for the *k*^th^ iteration of MCMC using the 95% isopleth of the kernel function,

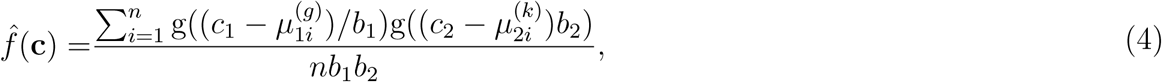

evaluated at locations of interest ***c*** *≡* (*c*_1_, *c*_2_) ′, choice of kernel function g(*·*), and bandwidth parameters *b*_1_ and *b*_2_. The result is a posterior distribution of the 95% home range isopleth, which could be used to further derive a posterior distribution of the home range area, thus fully incorporating all uncertainties in our estimate.

We fit the ATM and derived a convex hull and kernel density home range for individual Gunnison sage-grouse for different seasons (breeding and summer) across all years of available data. We compare these results with home range estimates using estimated locations from Lenth’s MLE, thus ignoring location uncertainty. Regardless of home range estimator, we found the spatial arrangement of the Gunnison sage-grouse home range was often different depending on whether location uncertainty was considered (Fig. 3c, Appendix S6). Ignoring location uncertainty often leads to overly small home range area estimates when compared to the estimate obtained when incorporating uncertainty. The contiguity of the kernel density home range was often affected by location uncertainty. Without taking into account location uncertainty, comparing home range area estimates across individuals could lead to highly biased inferences.

## Conclusion

Our model developments have important implications for interpreting historical radio-telemetry data analyses and to the future designs of these studies. While state-of-the-art tracking technologies (e.g., GPS) are increasingly used, animal telemetry via VHF radio is still widely used and will likely continue due to its low cost and miniaturization (Ponchon et al. 2013); digital VHF is increasingly used to study small-bodied migratory birds (Loring et al. 2017).

The development of the ATM addresses several complicating factors when dealing with azimuthal data. Foremost is that our model appropriately characterizes azimuthal telemetry uncertainty and allows this uncertainty to synthetically be propagated into spatial models. Appropriately accounting for uncertainties in ecological inference is needed to ensure appropriate inference (Brost et al. 2015; Hobbs and Hooten 2015; Figs. 3, 4). The ATM illustrates that the magnitude and shape of location uncertainty from azimuthal telemetry data is complex and highly variable. Previous methods have led to over confidence in the precision of animal locations, the certainty in resource selection, and the size of home ranges.

The ATM overcomes the issue of limited experimental field trials by allowing telemetry uncertainty to be directly modeled, thus accounting for telemetry uncertainty in location estimates. If the goal is to minimize location uncertainty, we found that it is prudent to encircle the animal, as well as obtain more than three azimuths (Fig. 4d, Table 1, Appendix S2: Fig. 3). However, the optimal study design will ultimately depend on the questions being considered (e.g., home range vs RSF study); researchers can pair the ATM with spatial models to identify optimal study designs that minimize logistical costs and maximizing model performance, something that was not previously possible.

We found the effects of location uncertainty on ecological inference is not straight-forward. Our RSF investigation demonstrated how location uncertainty affect on parameter estimates depends on the definition of availability (Hooten et al. 2013), whether covariates were categorical or continuous, and the degree of spatial autocorrelation in the covariate. Our simulation clarified that incorporating location uncertainty helps reduce bias in RSF coefficients across all levels of covariate spatial autocorrelation. Furthermore, our home range results suggest that previous studies that ignored location uncertainty could have been overly conservative in their estimate of home range areas; ignoring location uncertainty can have pernicious effects in terms of the shape and size of home range estimates.

## Acknowledgments

Funding was provided by CPW 1701 and NSF DMS 1614392 awards. M. L. Phillips led the Gunnison sage-grouse field data collection effort. Any use of trade, firm, or product names is for descriptive purposes only and does not imply endorsement by the U.S. Government. The findings and conclusions of the U.S. Fish and Wildlife Service employees in this article are their own and do not necessarily represent the views of the U.S. Fish and Wildlife Service.

